# Neural network models of the tactile system develop first-order units with spatially complex receptive fields

**DOI:** 10.1101/164954

**Authors:** Charlie W. Zhao, Mark J. Daley, J. Andrew Pruszynski

## Abstract

First-order tactile neurons have spatially complex receptive fields. Here we use machine learning tools to show that such complexity arises for a wide range of training sets and network architectures, and benefits network performance, especially on more difficult tasks and in the presence of noise. Our work suggests that spatially complex receptive fields are normatively good given the biological constraints of the tactile periphery.

## Results

First-order tactile neurons in the glabrous skin of the human hand have distal axons that branch in the skin and form many transduction sites^1-3^, yielding spatially complex receptive fields with many highly sensitive zones^4,5^ (**Fig. 1a**). We have recently shown that this arrangement permits first-order tactile neurons to signal high-level features of touched objects such as the orientation of a touched edge^4^, a capacity previously considered a hallmark of processing in the somatosensory cortex^6-8^. Here we leverage machine learning tools to investigate why complex receptive fields arise and what computational benefits they yield. We show that complex receptive fields arise under a wide range of training sets and biologically realistic network constraints. We also show that complex receptive fields benefit network performance, especially on more complex discrimination tasks and in the presence of noise.

**Figure 1.**
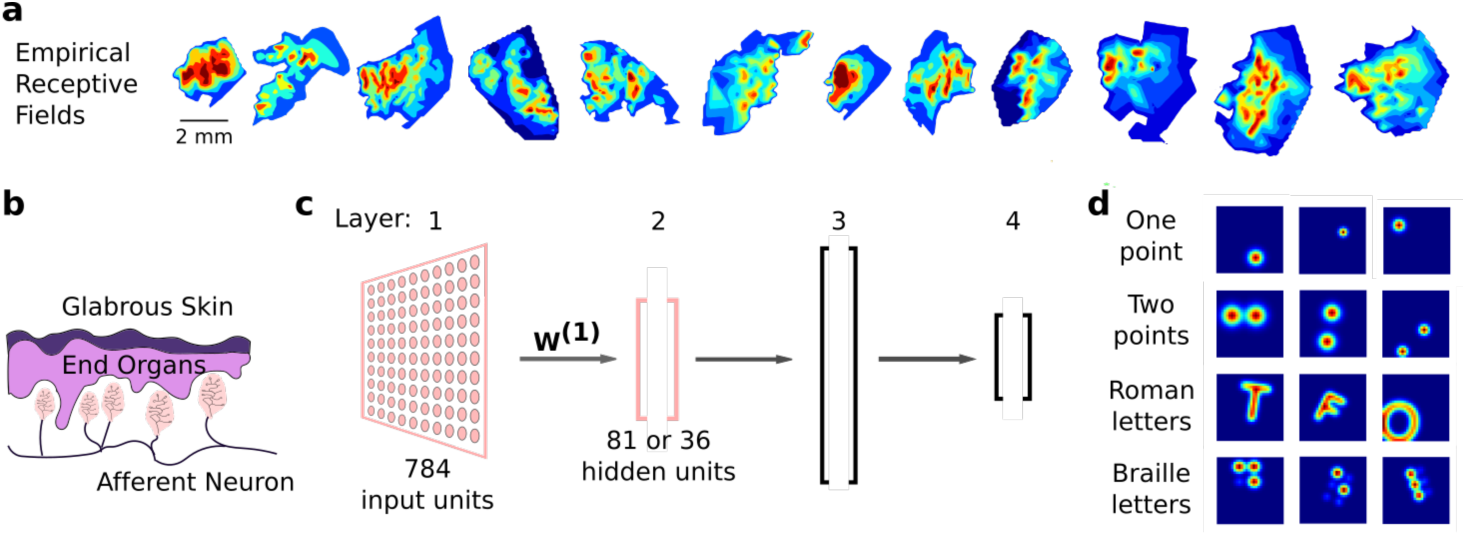
Theoretical and analytical setup. **(a)** Examples of receptive fields from human first-order tactile neurons terminating in the fingertip acquired via microneurography. Color indicates the relative firing rate of the neuron when stimulated with a small punctate stimulus. For full details, see Pruszynski and Johansson (2014). **(b)** Graphic representation of a cross-section through the human glabrous skin. Note how a single afferent neurons branches and innervates multiple mechanoreceptive end organs. **(c)** Our four-layer feedforward neural network. The first layer models a small patch of skin, W^(1)^ represents receptive fields, and the second layer models first order neurons. Layers 3 and 4 are a functional abstraction of the central nervous system. The relative sizes of each layer are shown but not to scale. Arrows represent fully connected feedforward weights between subsequent layers. End organs and first order neurons in panel **(b)** are colour matched with the layers that represent them in the model. **(d)** Examples of training data used to represent tactile stimuli. Each stimulus is shown on a 28 x 28 step grid. Stimuli were passed through a Gaussian filter and randomly rotated and translated. Points data were also randomly scaled.

We abstracted the tactile processing pathway with a four-layer feedforward neural network (**Fig. 1b, c**). The input layer of our network consisted of 784 units, representing mechanoreceptors distributed over a small patch of skin. In this arrangement, the weight matrix between the input and first hidden layer - which we call *W*^(1)^ - represents the receptive fields of first-order tactile neurons. Our network was trained on a range of stimuli including single points, multiple points, as well as Roman and Braille characters (**Fig. 1d**). These stimuli were subjected to translation and rotation and were spatially filtered to crudely approximate skin mechanics. Importantly, we introduced three biologically-inspired constraints. First, non-negative regularization in *W*^(1)^ to simulate the fact that first-order tactile neurons can only be excited when their transduction sites are stimulated^9^. Second, convergence from the input to the first hidden layer to simulate the many-to-one convergence from mechanoreceptors in the skin to first-order tactile neurons traveling in the nerve^1-3^. Third, two distinct unsupervised and supervised training phases, representing the encoding and interpreting aspects of the tactile processing pathway, respectively.

We first asked under what conditions, if any, our network learns spatially complex receptive fields. In our main analysis, the 784 units in the input layer converged to 81 units in the first hidden layer, estimating the fact that first-order tactile neurons innervate on the order of ten mechanoreceptors^1-3^. We reasoned that the complexity of the training set would influence the complexity of the receptive fields^10^. We tested this idea with four training sets: Gaussian single points, mixed one and two Gaussian-points, Roman letters, and a mixed set that included one and two Gaussian points, Roman letters and Braille characters in equal proportions (see Methods). These training sets represent different degrees of structural complexity, and consist of stimuli that have been used in tactile studies in both human and animal models^11-15^ but were not meant to represent the natural statistics of tactile stimuli, which are unknown.

We trained our network on each of these training sets in an unsupervised fashion and examined the resulting receptive fields (i.e. the *W*^(1)^ matrix). All networks, even those trained with the simplest training set, exhibited receptive fields with multiple areas of high sensitivity (**Fig. 2a**). Overall, there was a clear effect of training set on receptive field complexity (F(3, 76) = 1642, P<0.01) where the number of highly sensitive zones increased with the complexity of the training set (**Fig. 2b**). A similar effect was evident when analyzing receptive fields in the spatial frequency domain, with more complex training sets yielding higher spatial frequency content.

**Figure 2.**
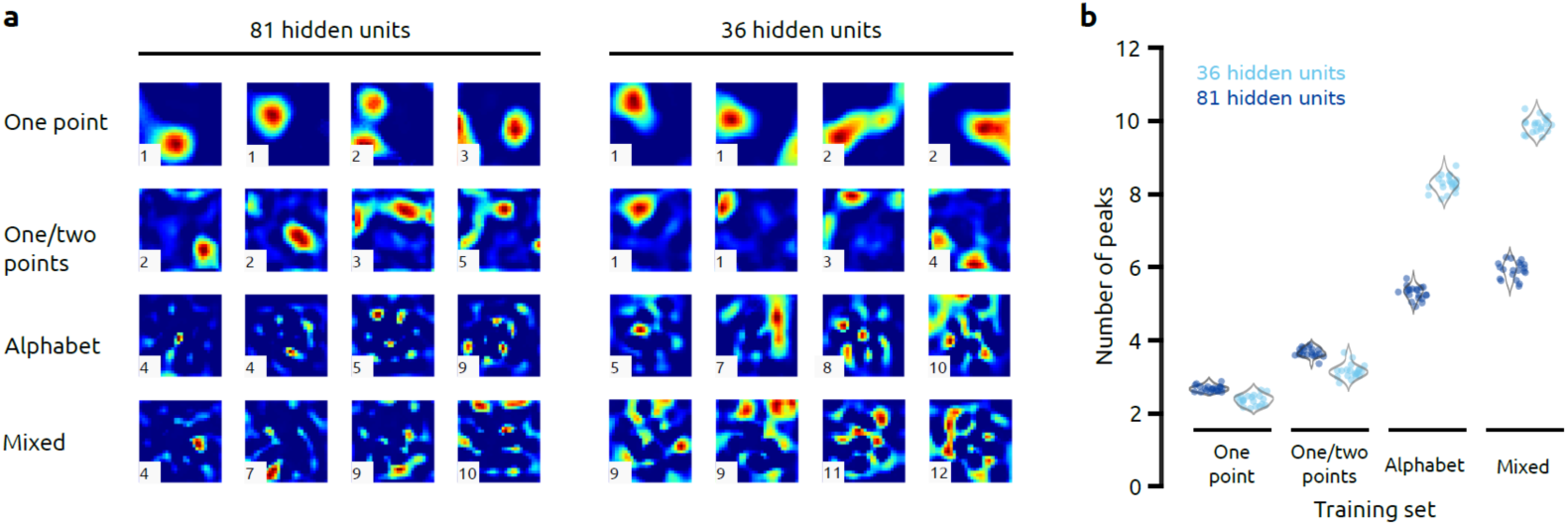
Analysis of receptive fields. **(a)** Examples of receptive fields learned by the 81- and 36-hidden unit models after training on different training sets (rows). Each receptive field is shown on a 28 x 28 step grid. Heat maps show areas with high weight values, which represent highly sensitive zones. Samples were chosen to show a variety of receptive field morphologies. The number on the bottom left corner of each receptive field is the number of peaks returned by our peak counting algorithm, which measures receptive field complexity. **(b)** The average complexity of each network under different architectures and training sets. Each data point is the mean peak count of receptive fields from that model on one iteration, with grey violin plots showing the overall frequency distribution across the 20 iterations we performed for each architecture and training set.

We next asked how the degree of convergence between the input and first hidden layers influenced receptive fields. That is, how physical constraints placed on the number of first-order tactile neuron axons traveling within the peripheral nerve should affect connectivity to mechanoreceptors in the skin. We reasoned that increasing convergence would increase receptive field complexity, since this smaller set of units must still encode the same set of inputs. We tested this idea by decreasing the size of the first hidden layer from 81 to 36 units, closer to the lower limit of biologically relevant convergence^1-3^, and training the network on the same four training sets described above. Increasing convergence did result in more complex receptive fields for alphabet and mixed networks (**Fig. 2b**). On average, the 36-unit alphabet network had 3.0 more peaks than the 81-unit alphabet network (t(38) = 46.39, P<0.01), and the 36-unit mixed network had 4.0 more peaks than the 81-unit mixed network (t(38) = 56.93, P<0.01). Interestingly, however, the one point and the one and two point networks (our simplest training sets) did not show increased complexity with increased convergence (**Fig. 2b**). In fact, the 36-unit one point network had 0.3 fewer peaks than the 81-unit one point network (t(38) = −8.55, P<0.01), and the 36-unit one and two point network had 0.5 fewer peaks than the 81-unit one point two point network (t(38) = −10.00, P<0.01).

At this point we further abstracted our network constraints to examine how they influenced the learned receptive fields. First, we trained our network on the mixed stimulus set without non-negative regularization in *W*^(1)^ and found qualitative changes in receptive field morphology such that they no longer had structural similarities to our previously documented empirical receptive fields^4^ (**Fig. 3a**). Second, we trained our network on the mixed stimulus set with extreme convergence (4 units in the first hidden layer) and, again, found the resulting receptive fields did not resemble our empirical receptive fields (**Fig. 3b**). Last, we trained our network on each of the four stimulus sets without convergence (i.e. 784 units in the first hidden layer). We reasoned that such a network may not develop complex receptive fields because it did not need to compress the input space, especially for the single dot training set given its simple spatial statistics. However, receptive fields with multiple highly sensitive zones emerged for all training sets to varying degrees (**Fig. 3c**).

**Figure 3.**
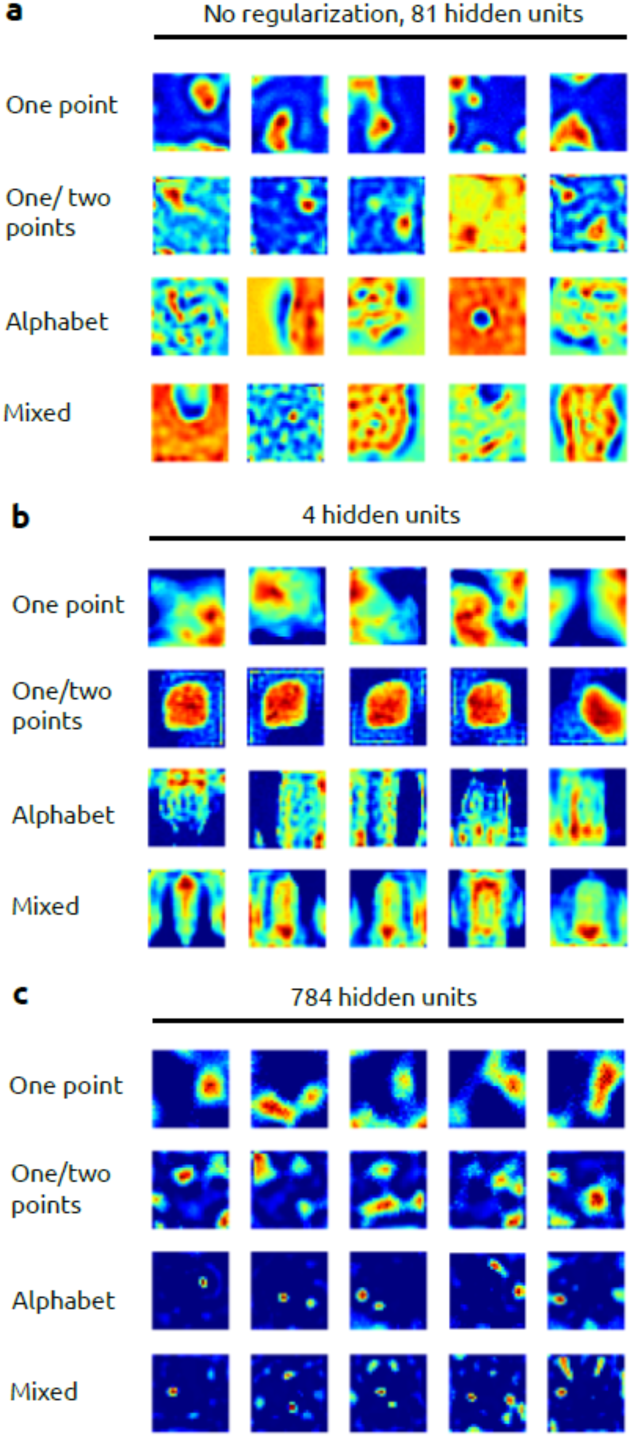
Alternative architectures. Same format as Figure 2 but showing exemplar receptive fields learned by three alternative networks featuring architectures with relaxed constraints.

Given that our networks developed complex receptive fields under all network constraints and training sets, we investigated the functional consequences that such an arrangement had on sensory processing. In these analyses, we trained the network on unlabelled Mixed stimuli, then fixed *W*^(1)^ and trained the remaining layers as a classifier using labelled Mixed stimuli. In our approach, the unsupervised training phase represents the encoding function of the tactile processing pathway, while the supervised training phase abstracts the more interpretive functions of the central nervous system. We compared this learned network against a network engineered to have single-peaked Gaussian receptive fields in *W*^(1)^ on discrimination and identification tasks. For the engineered network, we selected the width of the Gaussian receptive field (SD = 3.0 steps) that resulted in best performance.

We first asked whether complex receptive fields benefit spatial accuracy. We had the network perform two-point discrimination, a task central to many studies of tactile acuity^11,16,17^. Specifically, we used a two-alternative forced choice paradigm and defined the difference limen as the separation distance between stimuli at which the network classified 75% of the stimuli correctly. The learned network had a mean difference limen of 6.94 (SD = 1.36) steps on our input space, which corresponds to a modelled distance of ~1-3 mm, depending on assumptions about mechanoreceptor innervation density. Overall, performance of learned and engineered networks were not significantly different with 81 units in the first hidden layer (t(45) = −1.85, P = 0.071; **Fig. 4a**). Moreover, changing the degree of convergence from 81 to 36 units did not cause a statistically significant change in performance for either the learned or the engineered network (F(1, 82) = 0.31, P = 0.58; **Fig. 4a**).

**Figure 4.**
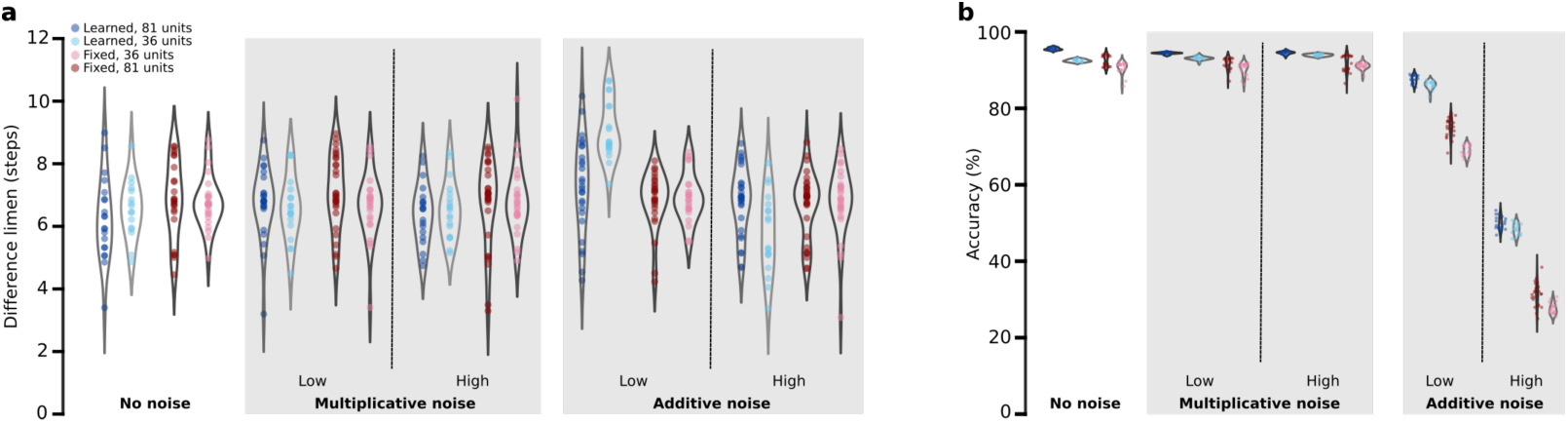
Model performance. Performance of 81- and 36-hidden unit models either trained on mixed stimuli or engineered with fixed Gaussian receptive fields on the **(a)** two-point discrimination and **(b)** alphabet classification tasks. **(a)** Data points show the difference limen, defined as the separation distance at which the model classifies 75% of 2000 test points correctly. **(b)** Data points show the overall classification accuracy of 7800 tested Roman letters. Grey violin plots show the frequency distribution of difference limens and accuracy across model iterations. Performance is reported at varying levels of multiplicative or additive noise (see Methods). Groups may have different numbers of data points as some networks failed to converge and were not considered for testing.

We then asked whether complex receptive fields benefit network performance in a more difficult identification task. We assessed the network’s ability to correctly classify characters from the Roman alphabet, as has been previously done with human participants^12^. In this case, engineering *W*^(1)^ to have single-peaked Gaussian receptive fields and increasing convergence both decreased network accuracy (F(1,79) = 103.78, P < 0.01, F(1, 39) = 107.23, P < 0.01, respectively), and the interaction between these factors was also significant (F(1, 79) = 7.05, P = 0.0096). That is, both learned and engineered networks performed well, but the learned networks outperformed engineered networks for both levels of convergence and the benefit of complex receptive fields increased with increased convergence (**Fig. 4b**).

Finally, we asked whether complex receptive fields benefit network performance in the presence of noise. We introduced varying levels of normally distributed additive and multiplicative noise to the training data during both unsupervised and supervised training phases and then tested the network’s performance on a noiseless dataset. The effect of training noise on the network’s ability to classify characters from the Roman alphabet was substantial (**Fig. 4b**). The learned network had an accuracy of 87.7% (SD = 1.1) with low levels of additive noise (see Methods) compared to 75.1% (SD = 2.5) for the fixed network with the same amount of noise, a statistically significant performance gap (t(41) = 20.65, P < 0.01). Convergence also significantly influenced classification accuracy under the different noise levels (F(6, 555) = 12.36, P < 0.01). The performance of the 36-unit network decreased by 1.4% compared to the 81-unit learned network with low levels of additive noise (t(38) = 4.25, P = 0.00013). In contrast, the performance of the 36-unit network with engineered Gaussian receptive fields decreased by 6.1% compared to the 81-unit engineered network (t(41) = 9.59, P < 0.01). The performance gap grew between learned and engineered networks with additional additive noise (**Fig 4b**). For all networks, multiplicative noise had a similar effect but much smaller effect size (**Fig. 4b**).

## Discussion

A core feature of the tactile processing pathway is that there are many more mechanoreceptors in the skin of the hand than there are first order tactile neurons in the median and ulnar nerves. It is not surprising, therefore, that first order tactile neurons branch^1-3^ since this is the only way they can innervate all the available mechanoreceptors. What may be surprising is the spatial complexity and apparent heterogeneity of the innervation pattern^4,5^, a feature which has been overlooked or ignored in previous models of the tactile processing pathway^11,18-20^. Our work here leverages simple machine learning tools to provide two fundamental insights in this respect. First, we show that spatially complex receptive fields are a normatively good and, perhaps, biologically parsimonious, arising under a wide range of training sets and network architectures. Second, we show that spatially complex receptive fields benefit network performance, especially in relatively difficult tasks and in the presence of noise.

Heterogeneously sampling the input space is a good thing for the nervous system to do because the input space of sensory stimuli is inherently sparse. Neural networks like the one we use here implicitly learn the statistical regularities (and thus sparsity) of the stimuli to which they are exposed. Indeed, such a machine learning approach has been shown to reproduce biological receptive field properties of neurons at various levels of the visual processing pathway^10,21^. Another suggestion for a mechanism to exploit sparsity comes from the field of compressed sensing, which shows that randomly sampling the input space can, under reasonable assumptions, allow a system to fully reconstruct a sparse input signal with fewer measurements than that prescribed by the Shannon-Nyquist theorem^22-25^. Given an input with sparsity *S* (at most *S* non-zero terms), in many situations the input signal can be fully reconstructed by randomly sampling at a frequency greater than 2*S* with no noise or multiplicative noise, or 4*S* with additive noise^22,24^, consistent with our observation that networks with more spatially complex receptive fields are particularly immune to additive noise. **Figure 5** illustrates a cartoon compressed sensing scenario in our experimental setting, showing that a network with fully randomized positive weights in the first hidden layer can perform strikingly well on the alphabet discrimination task relative to the learned and fixed networks we described above. That is, the random network performs only slightly worse than the learned network and equivalent to the fixed network with no noise and, as expected, is able to better maintain its performance as the amount of additive noise is increased. This is not to say that the heterogeneity of how first-order tactile neurons innervate mechanoreceptors is random – indeed random connectivity yields receptive fields that are qualitatively distinct from those we record from humans (**Fig. 5b**) – but, rather, that even random sampling can outperform pixel-like sampling with Gaussian receptive fields.

**Figure 5.**
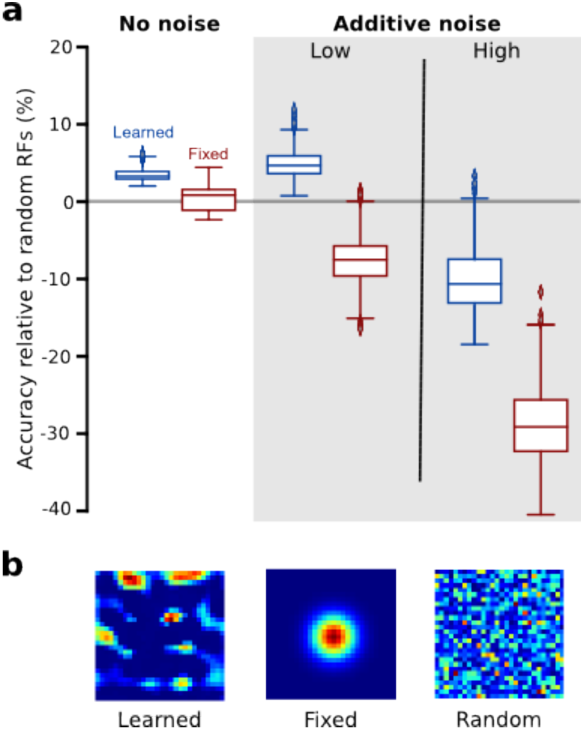
Comparison to compressed sensing framework. **(a)** Alphabet classification performance as a function of additive noise (same methodological details as in **Fig. 4b**) for the 81-unit learned and fixed models, relative to a network with the same architecture but random connectivity in the first hidden layer (n = 20 for each group). Box plot represents the first and third quartiles; whiskers extend to the 95^th^ percentile. **(b)** Example receptive fields from one representative unit in the learned, fixed, and the random models, respectively.

## Methods

### Feedforward Neural Network Architecture

We designed a four layer feedforward network model with layers *L*_1_ to *L*_2_) containing *s*_1_ to *s*_4_ units respectively. *s*_1_ = 784, *s*_2_ = 81 or 36, *s*_3_ = 784, and *s*_4_ = 26 *or* 2 depending on if the network is trained to perform alphabet classification or two-point discrimination. The general form of feedforward computation was as follows:

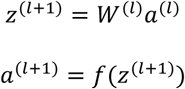

where *W*^(*l*)^ denotes the weights from layers *L*_*l*_ to *L*_*l*+1_, *z*^(*l*+1)^ is the weighted sum of outputs from layer *L*_*l*_, and *a*^(*l*)^ is the output of layer *L*_*l*_, after the activation function *ƒ*. For unsupervised training (*L*_1_ to *L*_3_), we used a rectified linear function *ƒ*(*x*) = *max*(0, *x*) for *W*^(1)^ and a softmax function for *W*^(2)^. For supervised training (*L*_1_ to *L*_4_), we used a rectifier for *W*^(1)^ and *W*^(2)^ and softmax for *W*^(3)^.

### Two-Phased Training and Non-Negativity Constraint

We randomly initiated weights by drawing from distribution *N*(0, 0.01). The general learning algorithm was mini-batch gradient descent with mini-batches of size 256. We trained the network in two phases. In the unsupervised learning phase, we trained *L*_1_ to *L*_3_ as an autoencoder that reproduced the input. The goal of gradient descent was to minimize the categorical cross-entropy cost:

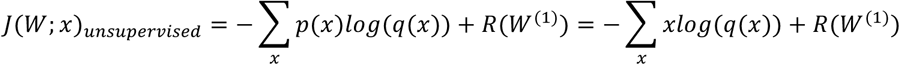

where, for training instance *x*, *p*(*x*) is the true output (which is equivalent to input *x* in the unsupervised learning phase), *q*(*x*) is the predicted input, and *R*(*W*^(1)^) is the non-negativity constraint, leading to the learning rule

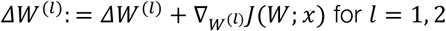

We incorporated the asymmetric regularization term^26^, *R*(*W*^(1)^), where

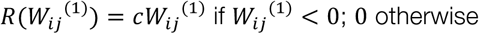

for each unit *j* of *L*_1_ and unit *i* of *L*_2_). *c* denotes an arbitrarily large constant, which we picked as 1000, that harshly penalized the network for learning negative weights in *W*^(1)^.

In the supervised phase, we froze *W*^(1)^ and trained *L*_1_ to *L*_4_ as a classifier. We reinitiated *W*^(2)^ between the two training phases. Depending on the discrimination task to be performed, the network may operate as a binary (for two-point discrimination) or multiclass (for alphabet) classifier. Gradient descent minimized the cross-entropy cost:

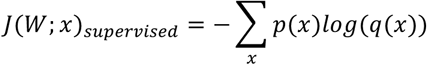

The learning rule in this phase was:

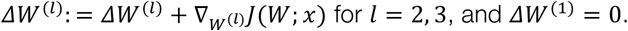

Network hyperparameters used during training varied among different network architectures and training sets. Networks that did not reach convergence in the number of iterations were removed from testing.

### Training Stimuli

We generated all training inputs *X* such that *x*_*ij*_ ∈ ℝ^28×28^. We generated Gaussian-points stimuli by initializing one or two peaks |*x*_*ij*_ | = 10 where *i*, *j* are integers chosen independently from distribution *U*(0,27), then passed through a two-dimensional Gaussian filter with width *σ* = 3.0.

We generated Roman letters stimuli as Helvetica characters normalized to 17 steps in height. We used similar height scaling for Braille characters. The filled portions of characters were initiated as |*x*_*ij*_ | = 1. We subjected each character to a random rotational angle drawn from distribution *N*(0, 20) in degrees, followed by random horizontal and vertical translation drawn from distribution *N*(0, 5) in steps.

We generated 60,000 training stimuli of each class. For Roman letters and Braille characters, there was approximately equal proportion of each character. Gaussian-points were evenly split between one and two points (i.e. 30,000 of each). We used standard one-hot encoding for labelling in supervised training.

### Receptive Field Complexity

We bootstrapped 1000 receptive fields from each network. First, we designed a peak counting algorithm that calculated the number of significant local maxima contained in each receptive field. For each receptive field *R*, we define *r*_*ij*_ as a peak if 1) it is a local maximum 2) |*r*_*ij*_| > (*max*_*k*_*r*_*k*_)/2, that is, the value of *r*_*ij*_ is greater than half of the global maximum, and 3) *r*_*ij*_ is at least 5 steps away from the next closest local maximum. These criteria prevent low amplitude noise from being counted as peaks. Second, we analyzed receptive fields in the frequency domain by performing discrete two-dimensional Fourier transformation using the Fast Fourier Transform algorithm. We performed Fourier transformation after normalizing sampled RFs by their peak values such that *max*_*k*_*r*_*k*_ = 1.0. Last, to compare information shared by each pair of networks, we used mutual information between pairs of bootstrapped RFs normalized by their respective entropies, such that 1.0 means perfect correlation and 0 means no mutual information. We binned weights into 10,000 bins before calculating mutual information so that the control group (learned versus learned) RFs has a normalized mutual information of close to 1.0.

### Model Performance

We assessed network accuracy in two-point discrimination and alphabet classification. We implemented two-point discrimination using a two-alternative forced choice paradigm. We generated 2000 one and two Gaussian-points in equal proportions. Two Gaussian-points were spaced symmetrically about the center of the input space at distances 0 to 22 steps apart with increments of 2 steps. We subjected two Gaussian-points to a random integer rotational angle drawn from distribution *U*(0, 90) in degrees. We defined the difference limen, or just-noticeable difference, for two-point discrimination as the distance at which the network correctly classified 75% of test stimuli. We estimated difference limen using cubic spline interpolation on the full accuracy plot.

We assessed the network on alphabet classification by testing it on 7800 new characters with 300 instances of each letter, subjected to rotational and translational variability as described above.

To assess robustness against noise, we trained the networks, in both unsupervised and supervised phases, with noisy data before testing them on noiseless data. We implemented multiplicative noise on input *X* as *ε*_*ij*_ = *c* · *u* · *x*_*ij*_ for each coordinate *i*, *j* in *X*, where *u* was randomly drawn from distribution *N*(0, 0.01). We implemented additive noise as *θ* = *c* · *v* · *max*_*k*_*x*_*k*_, where *v* was randomly drawn from distribution *N*(0, 0.01). We designated *c* = 1.0 as low-level noise and *c* = 3.0 as high-level noise. Noise was re-instantiated at the beginning of each training epoch.

## Acknowledgements

This work was supported by the Canadian Institutes of Health Research (Foundation Grant to JAP: 3531979). JAP received a salary award from the Canada Research Chairs Program.

## References

1. Cauna, N. Nerve supply and nerve endings in Meissner’s corpuscles. Am. J. Anat. 99, 315–50 (1956).

2. Cauna, N. The mode of termination of the sensory nerves and its significance. J. Comp. Neurol. 113, 169–209 (1959).

3. Nolano, M. et al. Quantification of myelinated endings and mechanoreceptors in human digital skin. Ann. Neurol. 54, 197–205 (2003).

4. Pruszynski, A. J. & Johansson, R. S. Edge-orientation processing in first-order tactile neurons. Nat. Neurosci. (2014). doi:10.1038/nn.3804

5. Johansson, R. S. Tactile sensibility in the human hand: Receptive field characteristics of mechanoreceptive units in the glabrous skin area. J. Physiol. 281, 101–123 (1978).

6. Bensmaia, S. J., Denchev, P. V., Dammann, J. F., Craig, J. C. & Hsiao, S. S. The representation of stimulus orientation in the early stages of somatosensory processing. J. Neurosci. 28, 776–786 (2008).

7. Yau, J. M. et al. Analogous intermediate shape coding in vision and touch. PNAS 106, 16457–62 (2009).

8. Fitzgerald, P. J., Lane, J. W., Thakur, P. H. & Hsiao, S. S. Receptive field properties of the macaque second somatosensory cortex: Representation of orientation on different finger pads. J. Neurosci. 26, 6473–84 (2006).

9. Grigg, P. Biophysical studies of mechanoreceptors. J. Appl. Physiol. 60, 1107–1115 (1986).

10. Olshausen, B. A. & Field, D. J. Emergence of simple-cell receptive field properties by learning a sparse code for natural images. Nature 381, 607–609 (1996).

11. Wheat, H. E., Goodwin, A. W. & Browning, A. S. Tactile resolution: Peripheral neural mechanisms underlying the human capacity to determine positions of objects contacting the fingerpad. J. Neurosci. 75, 5582–5595 (1995).

12. Vega-Bermudez, F., Johnson, K. & Hsiao, S. S. Human tactile pattern recognition: Active versus passive touch, velocity effects, and patterns of confusion. J. Neurophysiol. 65, 531–46 (1991).

13. Phillips, J. R., Johnson, K. O. & Hsiao, S. S. Spatial pattern representation and transformation in monkey somatosensory cortex. Proc. Natl. Acad. Sci. U. S. A. 85, 1317–21 (1988).

14. Johnson, K. O. & Lamb, G. D. Neural mechanisms of spatial tactile discrimination: Neural patterns evoked by braille-like dot patterns in the monkey. J. Physiol. 310, 117–144 (1981).

15. Phillips, J. R., Johansson, R. S. & Johnson, K. O. Representation of braille characters in human nerve fibres. Exp. Brain Res. 81, 589–592 (1990).

16. Johnson, K. O. & Phillips, J. R. Tactile spatial resolution. I. Two-point discrimination, gap detection, grating resolution, and letter recognition. J. Neurophysiol. 46, (1981).

17. Tong, J. et al. Two-point orientation discrimination versus the traditional two-point test for tactile spatial acuity assessment. Front. Hum. Neurosci. 7, 579 (2013).

18. Friedman, R. M., Khalsa, P. S., Greenquist, K. W. & LaMotte, R. H. Neural coding of the location and direction of a moving object by a spatially distributed population of mechanoreceptors. J. Neurosci. 22, 9556–9566 (2002).

19. Dodson, M. J., Goodwin, A. W., Browning, A. S. & Gehring, H. M. Peripheral neural mechanisms determining the orientation of cylinders grasped by the digits. J. Neurosci. 18, 521–530 (1998).

20. Saal, H. P., Delhaye, B. P., Rayhaun, B. C. & Bensmaia, S. J. Simulating tactile signals from the whole hand with millisecond precision. Proc. Natl. Acad. Sci. U. S. A. 201704856 (2017). doi:10.1073/pnas.1704856114

21. Yamins, D. L. K. et al. Performance-optimized hierarchical models predict neural responses in higher visual cortex. Proc. Natl. Acad. Sci. 111, 8619–8624 (2014).

22. Candes, E. J. & Wakin, M. B. An introduction to compressive sampling. IEEE Signal Process. Mag. 25, 21–30 (2008).

23. Candès, E. J., Romberg, J. & Tao, T. Robust uncertainty principles: Exact signal reconstruction from highly incomplete frequency information. IEEE Trans. Inf. Theory 52, 489–509 (2006).

24. Candès, E. J. The restricted isometry property and its implications for compressed sensing. Comptes Rendus Math. 346, 589–592 (2008).

25. Donoho, D. L. & L., D. Compressed sensing. IEEE Trans. Inf. Theory 52, 1289–1306 (2006).

26. Lemme, A., Reinhart, R. F. & Steil, J. J. Efficient online learning of a non-negative sparse autoencoder. in European Symposium on Artificial Neural Networks (2010).

